# Gamma secretase inhibitors inhibit ovarian cancer development and enhance olaparib’s anti-ovarian cancer activity and its mechanism

**DOI:** 10.1101/2020.05.19.103853

**Authors:** Yao Xiaoxiao, Zheng Nan

**Affiliations:** Zhejiang Shuren University Shulan International Medical College, Zhejiang Hangzhou, CHINA, 310000; Wenzhou Medical University, Zhejiang Wenzhou, CHINA, 325035

**Keywords:** ovarian cancer, olaparib, apoptosis, γ-secretase inhibitor, chemotherapy resistance

## Abstract

**Background:** Ovarian cancer is the fifth leading cause of cancer-related deaths in women. Although cytoreductive surgery combined with chemotherapy and / or targeted therapy has achieved a certain effect, but advanced patients have limited clinical benefits and are prone to relapse. Among them, Notch / jagged1 and Wnt / β-catenin signal transduction plays an important role in the development of ovarian cancer and chemotherapy resistance, and it is very important to find new effective therapeutic drugs to inhibit tumor development and increase tumor chemotherapy sensitivity.

**Methods:** To establish ovarian cancer xenotransplantation model, to detect the effects of γ-secretase inhibitor, olaparib and combination drugs on the development of ovarian cancer, and to detect the proliferation and apoptosis of ovarian cancer cells with γ-secretase inhibitor, olaparib and combination drugs .After knocking down Jagged1 or β-catenin, the protein expression levels of related signaling pathways under different drug treatments were analyzed by immunoblotting, and related gene expression changes were analyzed.

**Results:** DAPT reduced β-catenin expression in a proteasome-synthetic pathway-dependent manner, thereby inhibiting the synthesis and transcription of Jagged1 protein, thereby inhibiting the expression of Notch signal transduction pathway-related proteins and gene transcription, inhibiting the activity of ovarian cancer cells and inducing cell apoptosis death, enhance the anti-ovarian cancer activity of olaparib.

**Conclusion:** DAPT inhibits cell proliferation, induces cell apoptosis, inhibits tumor development in vivo through β-catenin / Jagged1 inhibition, and exerts anti-ovarian cancer activity sensitizing effect of olaparib.

## 1 Introduction

Ovarian cancer is a common malignant tumor of the female reproductive system, and its death rate ranks first in female malignant tumors. Due to early metastasis, early diagnosis, and chemotherapy resistance, most patients are found to be late ^[1]^. Early ovarian cancer is mainly treated by various methods including surgical resection, chemotherapy, radiotherapy, and embolization, but advanced and unresectable ovarian cancer is extremely difficult to treat, which makes it difficult for existing treatments to meet clinical medical needs. The patient has a poor prognosis and mortality Very high ^[2–3]^. Olaparib was approved for marketing in China in September 2018. It is a PARP inhibitor targeted for the treatment of ovarian cancer. It is currently the only drug approved by the United States and the European Union for the treatment of patients with advanced ovarian cancer with BRCA mutations. It has gradually become the first-line drug for the treatment of advanced ovarian cancer ^[4–5]^, but due to the rapid development of acquired drug resistance and multi-drug resistance, the clinical treatment effect of olaparib is not good, and there is an urgent need for the development of ovarian cancer Of more effective therapies or synergists of olaparib.

Notch / Jagged signaling pathway is a very important signal transduction system for regulating cell proliferation, differentiation, apoptosis and other cell functions, including Notch receptors (Notch1, Notch2, Notch3, Notch4), Notch ligands (Delta-like1, 3, 4 and Jagged1,2) and other effectors. It is reported in the literature that the activation of Notch signal is closely related to the development of ovarian cancer and chemotherapy resistance, and can promote the progress of ovarian cancer through the interaction with various cytokines and signaling pathways ^[6–9]^. Therefore, by interfering with the upstream or downstream genes of Notch signal to inhibit the activation of Notch signal may become a potential mechanism for the treatment of ovarian cancer and enhance the sensitivity of ovarian cancer radiotherapy.

The γ-secretase complex is composed of four proteins: presenilin, nicastleline, APH-1 (prepharyngeal lesion factor 1) and PEN-2 (presenilin enhancer 2). γ-secretase complex can indirectly promote the release of Notch intracellular domain (NICD) and translocate into the nucleus to act as a translation factor recombinant hairless binding protein inhibitor (RBPSUH) (also known as RBP-Jκ or CSL) Co-activator, which activates Notch / Jagged signal transduction ^[10–11]^. RBP-Jκ is a DNA-binding component of the Notch signal transduction pathway. It can encode the Hes-related enhancer gene and genes related to the YRPW motif (Hey) family protein by activating the target gene, and many of them include ovarian cancer. Cancers are closely related ^[12–13]^.

DAPT (DAPT) is a γ-secretase substrate that can act as a γ-secretase inhibitor to indirectly inhibit Notch signal transduction in autoimmune and lymphoproliferative diseases including Alpine disease and lupus erythematosus (SLE) The medicinal effect is significant, and it also plays an important role in the growth of various cancer cells, angiogenesis and differentiation of human induced pluripotent stem cells (hIPSC) by inhibiting Notch signaling ^[14–15]^. The purpose of this study was to explore the effect of DAPT on hepatocellular carcinoma in vivo and in vitro, and the effect of DAPT on the anticancer activity of olaparib, and to preliminary explore the potential mechanism of DAPT’s antitumor effect.

## 2 Methods and reagents

### 2.1 Reagent

DAPT, proteasome inhibitor MG262, dimethyl sulfoxide (DMSO), GSK3β inhibitor TWS119 and SB216763 were purchased from Biolab Reagents (China); Wnt agonist WntC59, Glo lysis buffer were purchased from Biyuntian reagent Company (China), BCA protein determination kit was purchased from Kunshan Jincheng Reagent Company (China). Notch1, Notch2, Notch3, Jagged1, cleaved Notch1 (NICD), Hes1, β-catenin, phosphorylated β-catenin, c-Myc, cyclin D1, survivin and GAPDH monoclonal primary antibodies were purchased from Gloucester Biotechnology Corporation (China).

### 2.2 Cells and culture

SKOV3 and OVcar3 ovarian cancer cell lines were purchased from Shanghai Institute of TWS119chemistry and Cell TWS119logy, Chinese Academy of Sciences. Human ovarian cancer cell lines SKOV3 and OVcar3 cells were cultured in DMEM medium (HyClone Logan Company) containing 10% fetal bovine serum, 100 U· ml^−1^ penicillin and 100 μg· ml^−1^ streptomycin. Incubate in a constant temperature and humidity incubator.

### 2.3 Determination of cell viability

When the cells grow to 70% confluence, trypsinize and collect the cells, use DMEM medium to dilute the cells to 1×10^5^ · ml^−1^, inoculate the cells in a 96-well plate and culture overnight (3000 cells/well), then use DAPT, Olaparib or combination drugs were treated at the specified concentration for 24, 48 or 72 h, and then according to the reagent manufacturer’s instructions, add 10 μl of Cell Counting Kit-8 (CCK8) reagent (Dojindo, Japan) to each well. After incubating at room temperature for 2 h. The absorbance was measured at 450 nm and normalized to the background absorption of the culture medium in the absence of cells. The experiment was repeated three times.

### 2.4 Soft agar colony formation test and clone formation test

When the cells grow to the logarithmic growth phase, trypsin digestion to collect the cells, use DMEM medium to dilute the cells to 1×10^6^ · ml^−1^, inoculate the cells into a 6-well plate (1000 cells / well), after 24 hours of culture, Incubate for 10-14 days with 1% 0.33% BME agar medium with or without DAPT (0–1 μM) in 10% FBS^[15]^. Cell colonies were counted under the microscope using Image-Pro Plus software (Media Cybernetics, USA).

### 2.5 Flow cytometry analysis of apoptosis

Follow the manufacturer’s instructions to check the apoptosis with Annexin V-FITC Apoptosis Detection Kit (Becton Dickinson TWS119sciences, USA): After treating cells with DAPT, olaparib or combination drugs for 24 hours, Annexin V-FITC FITC and propidium iodide (PI) staining^[17]^, after incubation at room temperature for 5 minutes in the dark, the apoptosis was analyzed by flow cytometry (Becton Dickinson FACS Vantage SE, USA).

### 2.6 Transfection of cells

According to the manufacturer’s instructions, use lipofectamine 3000 (Invitrogen, USA) to transiently transfect sh Jagged1, shβ-catenin or pcDNA3 plasmids (negative control, si-NC) in Opti-MEM medium (Gibco, USA) into SKOV3 cells^[18]^ ,After 24 hours of transfection, cell viability and apoptosis were detected. In addition, the cells were treated with DMSO or DAPT (4 μM) for 4 hours and whole cells were extracted for Western blotting.

### 2.7 Protein preparation and Western blot analysis

When the cells grow to the logarithmic growth phase, trypsin digestion, 1000 r, centrifugation for 5 min to collect the cells, add RIPA buffer (Qingke TWS119technology Company, China) for lysis, and add 1x protease inhibitor (Roche Applied Sciences, Germany). According to the manufacturer’s instructions, use PierceTM BCA protein detection kit (Acbam, China) for protein quantification. At 120V, perform the same amount of protein on 10% SDS-polyacrylamide gel electrophoresis (PAGE) for 90 min, and then transfer to poly On a vinylidene fluoride (PVDF) membrane (TWS119-Rad, USA), the membrane was blocked in 5% skim milk at room temperature for 1 hour, then incubated with a specific primary antibody at 4℃ overnight, and then with horseradish Peroxidase-labeled goat anti-rabbit IRDye 800CW secondary antibody (Agilent TWS119technology, China) was incubated at room temperature for 2h. According to the manufacturer’s instructions, an Odyssey infrared imaging system (LI-COR, USA) was used to capture the image. The protein density was measured using Quantity One imaging software (TWS119-Rad, USA).

### 2.8 RNA extraction and real-time PCR

According to the manufacturer’s instructions, total RNA was extracted from ovarian cancer cell lines using the TRIzol kit (TaKaRa, Japan), and reverse transcription was performed using cDNA reverse transcription kit (Thermo Fisher, Germany). Follow the instructions to reverse transcribe cDNA. Design primers based on GenBank sequence. RT-qPCR amplification was performed using SYBR Premix Ex Taq TM II kit (TaKaRa Corporation, Japan). Table 1 shows the PCR primers used to amplify the indicated genes. The relative expression of the samples was normalized to the internal control GAPDH.

**Table 1.**
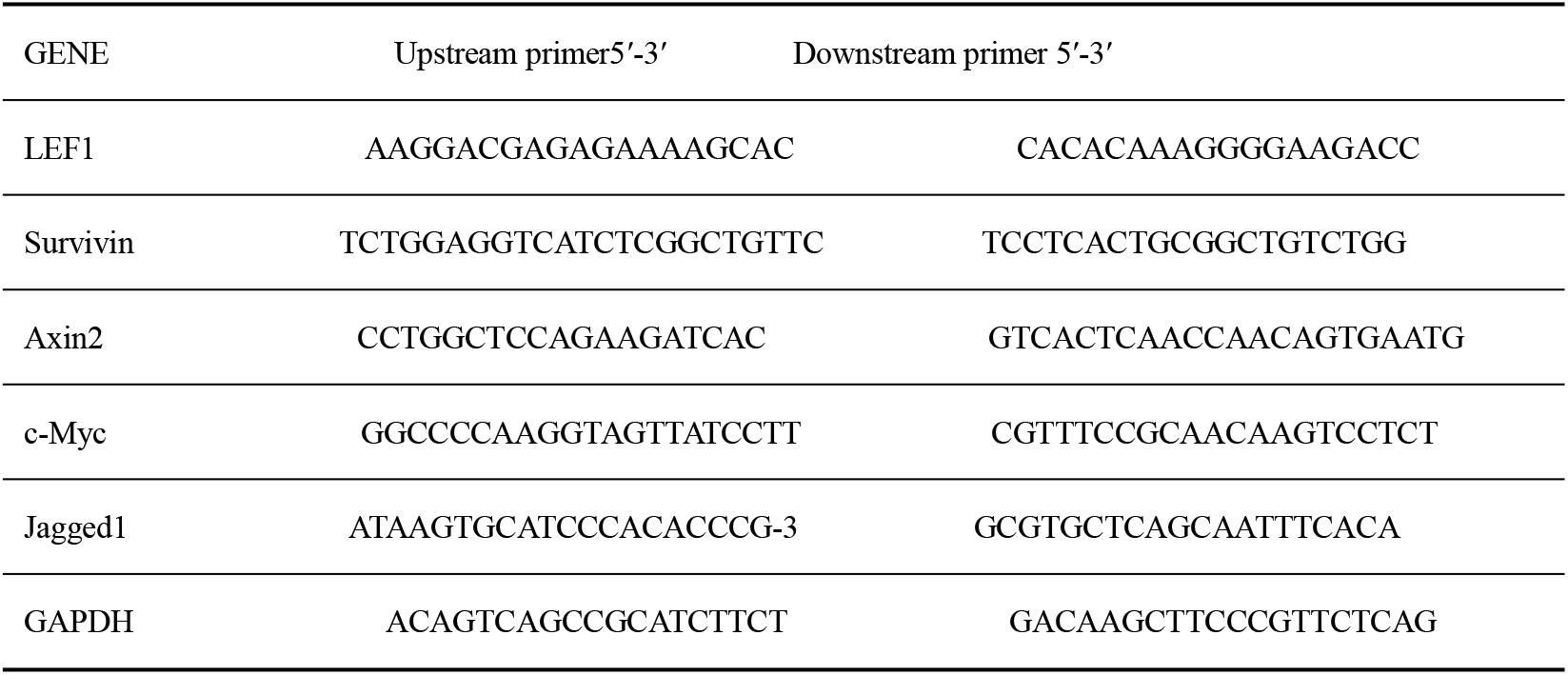
Primer sequences used in PCR

### 2.9 Immunocytochemistry (ICC) and immunohistochemistry (IHC)

Cell culture slides with a fusion degree of 70% were washed with PBS for 5 min × 3 times in the culture plate. 4% formaldehyde was fixed for 30 min, washed with PBS three times, permeabilized with 0.2% Triton X-100 at room temperature for 30 min, washed with PBS three times, blocked with 3% BSA at room temperature for 30 min, and then DAPT (0.5-1 μM) Treat for 24 hours, then incubate with β-catenin primary antibody (1: 100) overnight at 4℃. After washing three times with TBST, incubate with Alexa Fluor 594 labeled goat anti-rabbit IgG secondary antibody (1: 200) for 2 h at room temperature, after TBST washing three times, stain with DAPI stain (CST, USA) according to the reagent manufacturer’s instructions. Fluorescence imaging was captured with LAS AF Lite software (Leica Microsystems, Germany) and Leica TCS SP8 X microscope (Leica Microsystems, Germany)

After the end of the animal experiment, the mice were sacrificed by cervical dislocation, the liver tissue was aseptically separated, fixed in neutral 10% formalin buffer for 24h, embedded in paraffin, and then sliced, the tissue section was deparaffinized, and after 10 min of antigen recovery in citrate buffer at pH = 6.0, the sections were treated with methanol containing 3% H_2_O_2_ to quench the activity of endogenous tissue peroxidase, followed by 1% bovine serum albumin (BSA) Incubate to block non-specific binding. IHC staining with specific antibodies: Ki67 primary antibody (1: 500), TUNNEL (1: 500) and β-catenin primary antibody (1: 500, CST, USA), horseradish peroxidase after 24h incubation at room temperature the coupled secondary antibody was stained, and the slides were developed in diaminobenzidine, counterstained with hematoxylin and observed under an Olympus microscope.

### 2.10 Animals

Sixty four-week-old male Balb/c nude mice were obtained from the Experimental Animal Center of Wenzhou Medical University (China) and kept in the SPF laboratory of the Experimental Animal Center without specific pathogens. All animal experiments were conducted in accordance with the 《Guidelines for the Care and Use of Laboratory Animals》of the National Institutes of Health, and were approved by the Animal Experiment Ethics Committee of Wenzhou Medical University. 5 × 106 SKOV3 cells in 100 μl PBS were injected subcutaneously into the right abdomen of the mice. When the tumor volume reached 50 mm^3^, the mice were randomly divided into 6 groups (n = 10 for each group) and injected with 0.75 mg · kg^−1^ DAPT, 1.5 mg · kg^−1^ DAPT, 10 mg · kg^−1^ opatinib, 40 mg · kg^−1^ opatinib, combination (0.75 mg · kg^−1^ DAPT +10 mg · kg^−1^ opal Tini), or PBS as a control, injected intraperitoneally once a day for 10 consecutive days of treatment. Calculate the tumor volume according to the following formula:

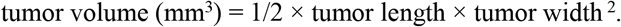

### 2.11 Statistical analysis

SPSS 25.0 software was used for statistical analysis. The measurement data is expressed by x ± s, and the statistically significant difference between the data is calculated using analysis of variance (ANOVA) or Student t test. All data are the results of at least 3 independent experiments. P <0.05 indicates that the difference is statistically significant.

## 3 results

### 3.1 DAPT inhibits ovarian cancer cell viability and induces apoptosis in vitro

In order to test the effect of γ-secretase inhibitor DAPT on the in vitro proliferation activity of ovarian cancer cells, we used different concentrations of DAPT (2, 4, 6, 8, 10 μM) to incubate with SKOV3 and OVcar3 ovarian cancer cell lines for 24 hours, and passed The CCK8 test was used to detect the activity of ovarian cancer cells, and the clonal proliferation test was used to detect the proliferation of ovarian cancer cells. The results show that DAPT can significantly inhibit the activity of ovarian cancer cells and inhibit the formation of SKOV3 and OVcar3 cancer cell colonies in a dose-dependent manner. In addition, flow cytometry analysis showed that DAPT treatment can significantly induce apoptosis of SKOV3 and OVcar3 cells at a dose Dependence (Figure 1D)

**Figure 1.**
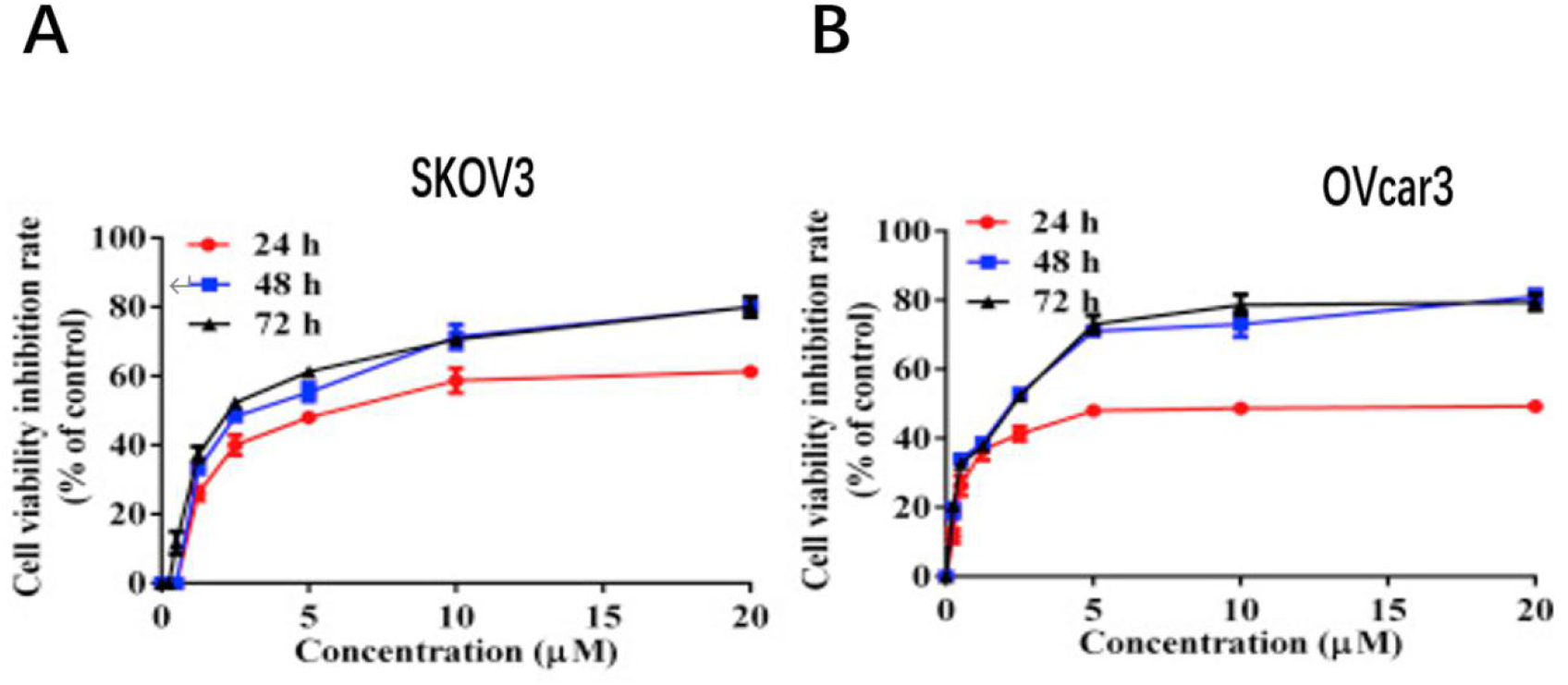
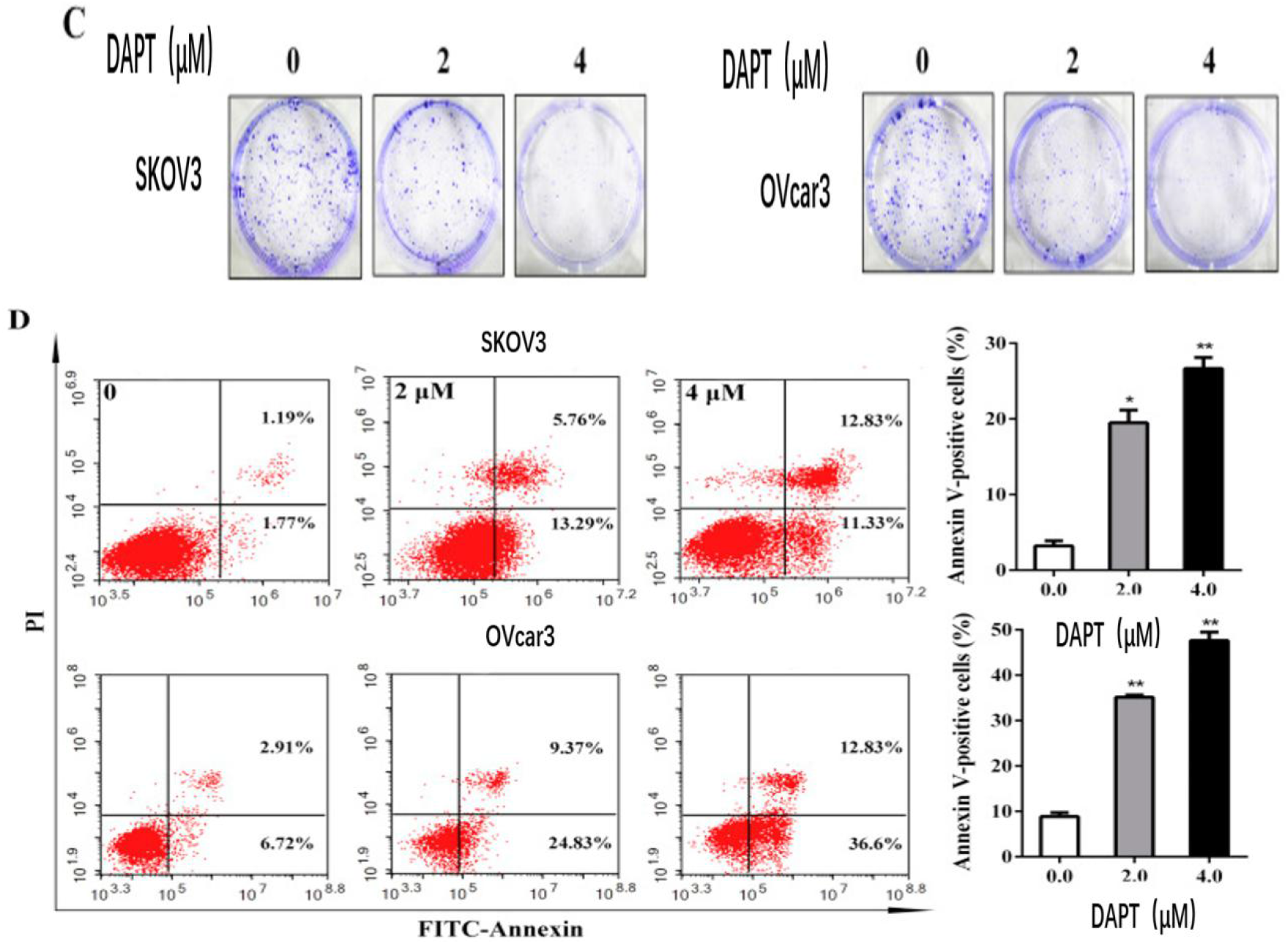
DAPT inhibits the activity of ovarian cancer cells. (A) Cell viability curves of human ovarian cancer SKOV3 and OVcar3 cells after DAPT treatment. (B) Activity curves of OVcar3 cells treated with DAPT at different doses / times (left),activity curves of SKOV3 cells treated with DAPT at different doses and times (right) (C) Treated with DAPT (2 μM, 4 μM) for 2 weeks representative images of colonies formed by OVcar3 and SKOV3 cells. (D) Representative flow cytometry apoptosis analysis graph (left) and quantification of apoptotic cells of OVcar3 cells (top) and SKOV3 cells (bottom) after DAPT (2μM, 4μM) treatment for 24 hours The data of the experiment repeated three times is expressed as mean ± SD, * P <0.05.

### 3.2 DAPT inhibits ovarian cancer tumor growth

In order to detect the antitumor activity of DAPT in vivo, we injected 5 × 10^6^ SKOV3 cells in 100 μl PBS subcutaneously into the right abdomen of the mice. When the tumor in the mice grew to about 50 mm3, the mice were randomly divided into 3 groups (n = 10), the control group mice were intraperitoneally injected with 5ml PBS, the low-dose group and high-dose group were injected with 0.75 mg · kg^−1^DAPT (dissolved in DMSO) and 1.5 mg · kg^−1^DAPT, once a day, continuous treatment 10 days. The results show that DAPT treatment can significantly inhibit the growth rate and volume of SKOV3 tumors (Figure 2A-D), and no weight loss was observed in mice.

**Figure 2.**
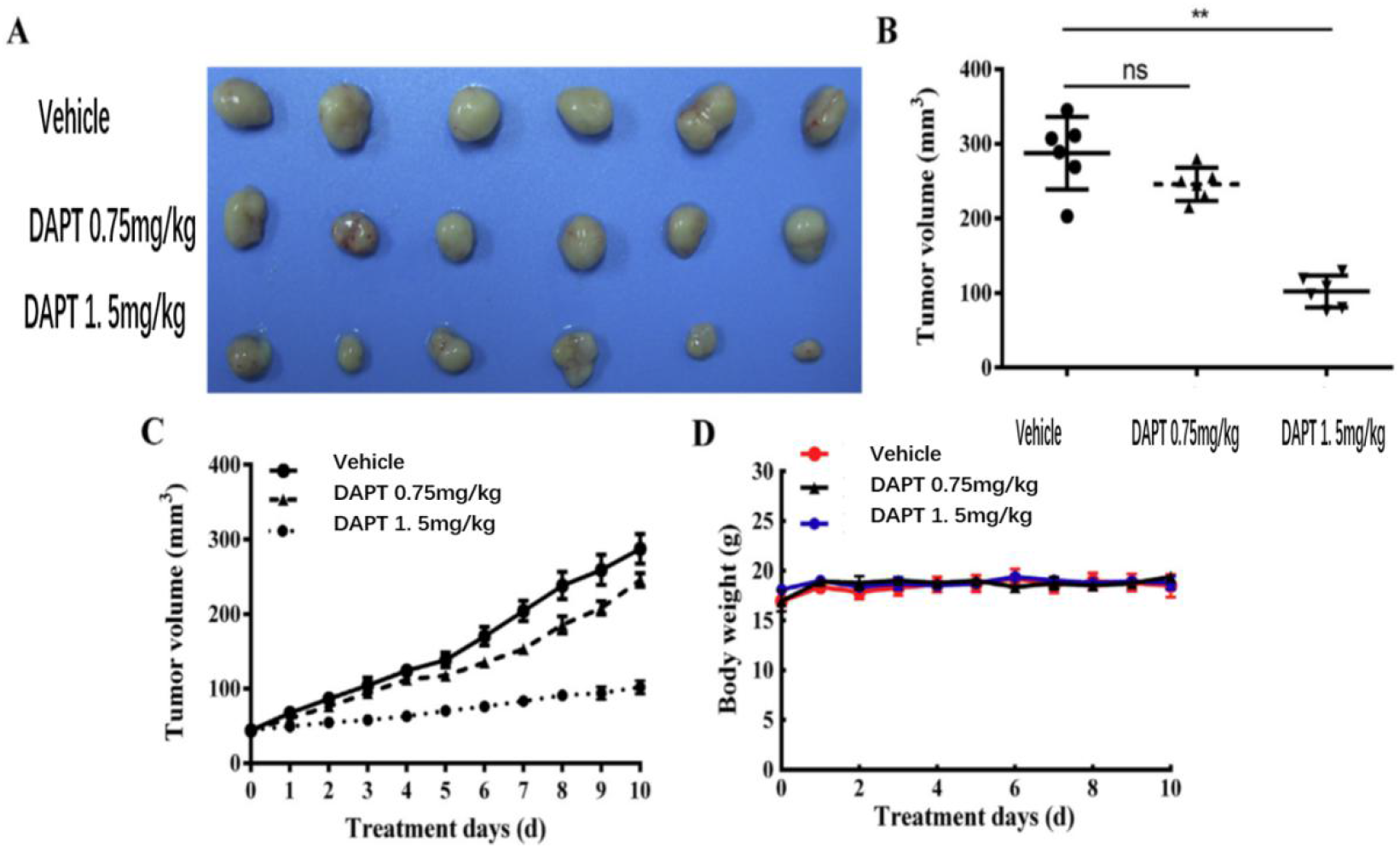
DAPT inhibits tumor growth in nude mouse models of ovarian cancer. Tumor images in a xenograft nude mouse model. (B) Tumor volume in a xenograft nude mouse model. (C) Tumor growth curve in a xenograft nude mouse model. (D) Body weight of mice during treatment. The data are expressed as mean ± SD. * p <0.05.

### 3.3 DAPT enhances the anti-ovarian cancer activity of olaparib in vitro

To test the effect of DAPT on the antitumor activity of olaparib in vitro, we used different concentrations of DAPT, different concentrations of olaparib, or a combination of drugs (4μM DAPT + 1.25μM olaparib) to treat the cells for 24 hours, and then Cell viability was detected by CCK8, and apoptosis was analyzed by flow cytometry. The results showed that after treatment with DAPT or olaparib cells, the activity of ovarian cancer cells was significantly inhibited, and showed a dose-dependent (Figure 3A, D), compared with DAPT or olaparib alone, the combination therapy can be significantly Inhibit cell viability, (Figure 3B, E, C, F). In addition, flow cytometry analysis showed that the combination of DAPT and olaparib can significantly induce the apoptosis of SKOV3 and OVcar3 cells compared with DAPT or olaparib treatment alone (Figure 3G and H), indicating that DAPT can be enhanced the anti-ovarian cancer activity of olaparib in vitro.

**Figure 3.**
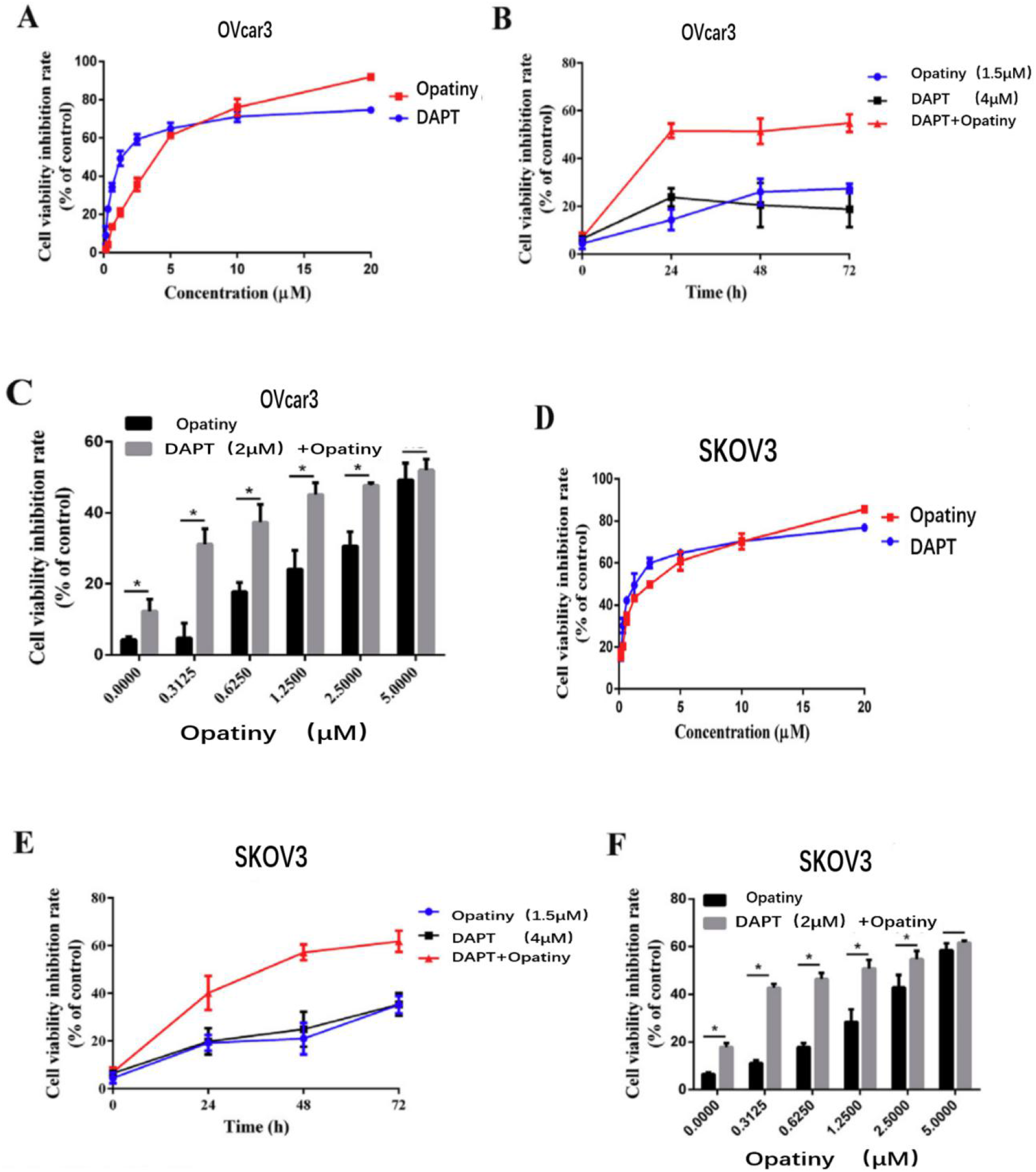
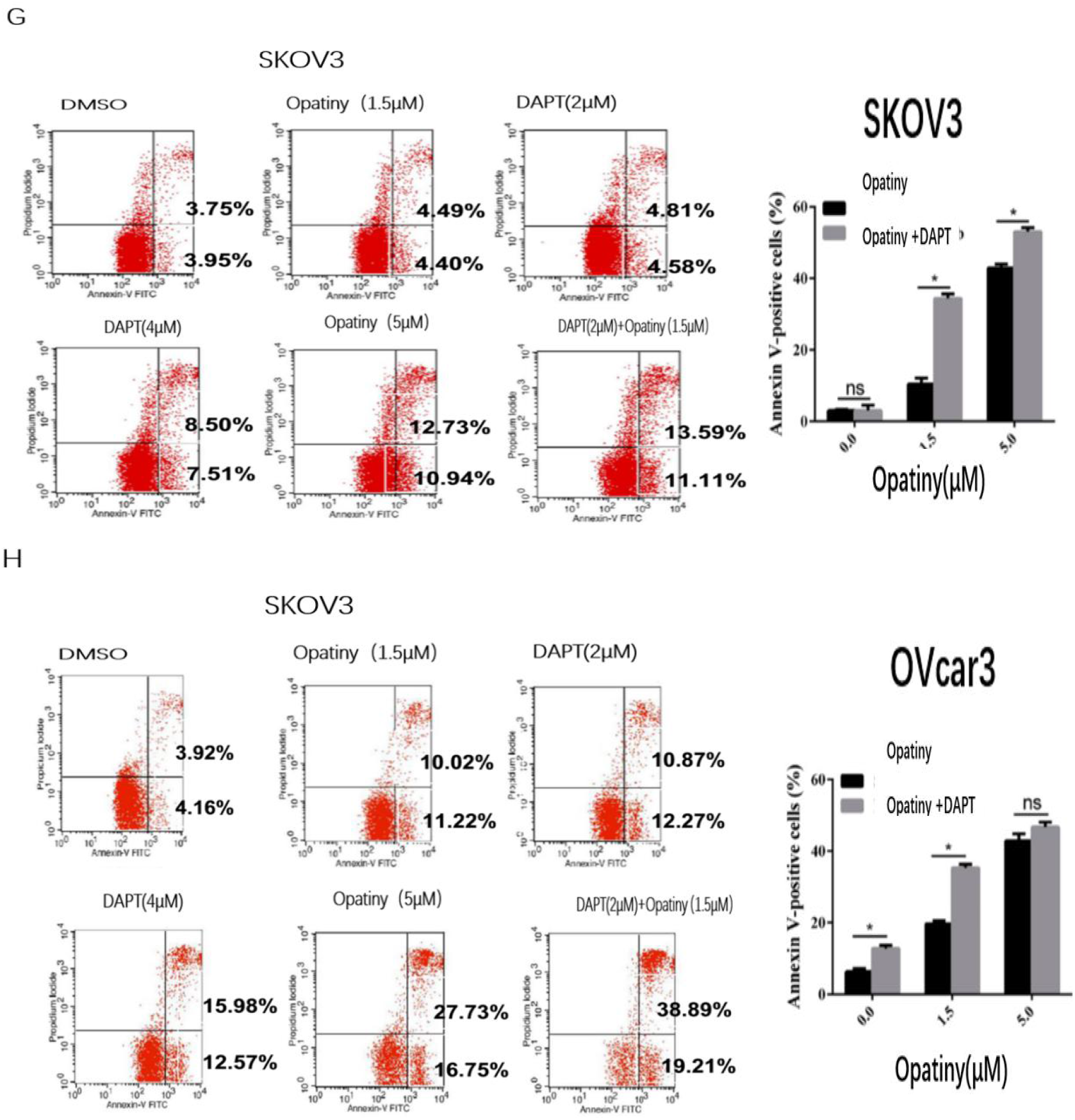
DAPT enhances the antitumor activity of olaparib in vitro. (A) The inhibition curve of DAPT and olaparib on human ovarian cancer OVcar3 cells. (B) Activity inhibition curves of DAPT, olaparib, and combined drugs on human ovarian cancer OVcar3 cells. (C) Quantitative inhibition of OVcar3 cell activity by olaparib treatment or combination therapy. (D) Activity inhibition curves of DAPT and olaparib on human ovarian cancer SKOV3 cells. (E) Activity inhibition curves of DAPT, olaparib, and combined drugs on human ovarian cancer SKOV3 cells. (F) Quantification of the inhibitory effect of olaparib treatment or combination therapy on SKOV3 cells (G) SKOV3 cell apoptosis and quantification after 24 hours of olaparib alone or combination therapy. (H) Apoptosis and quantification of OVcar3 cells 24 hours after olaparib alone or combination therapy. Data are expressed as mean ± SD, * P <0.05.

### 3.4 DAPT enhances the anti-ovarian cancer activity of olaparib in vivo

To test the effect of DAPT on the antitumor activity of olaparib in vivo, we injected 5 × 10^6^ SKOV3 cells in 100 μl PBS subcutaneously into the right abdomen of mice. And randomly divided into 5 groups (n = 6), control group mice were injected with 5ml PBS intraperitoneally, DAPT group was injected with 0.75 mg · kg^−1^DAPT (dissolved in PBS), olaparib low-dose group and high-dose group were injected 10 mg · kg^−1^ olaparib (dissolved in PBS) and 40 mg · kg^−1^olaparib, 0.75 mg · kg^−1^DAPT and mg · kg^−1^ olaparib were injected into the combination group, every day once injection, continuous treatment for 10 days. Ovarian cancer xenograft nude mice model results show that compared with the control group, DAPT, low/high dose olaparib, combined treatment can significantly inhibit tumor progression, and compared with DAPT (0.75 mg · kg^−1^) or low high-dose olaparib (10 mg · kg^−1^) treatment, the combined treatment can significantly inhibit tumor development, and it is equivalent to high-dose olaparib (30 mg · kg^−1^; Figure 4A) inhibition. It shows that DAPT can significantly enhance the anti-ovarian cancer activity of olaparib in vivo.

**Figure 4.**
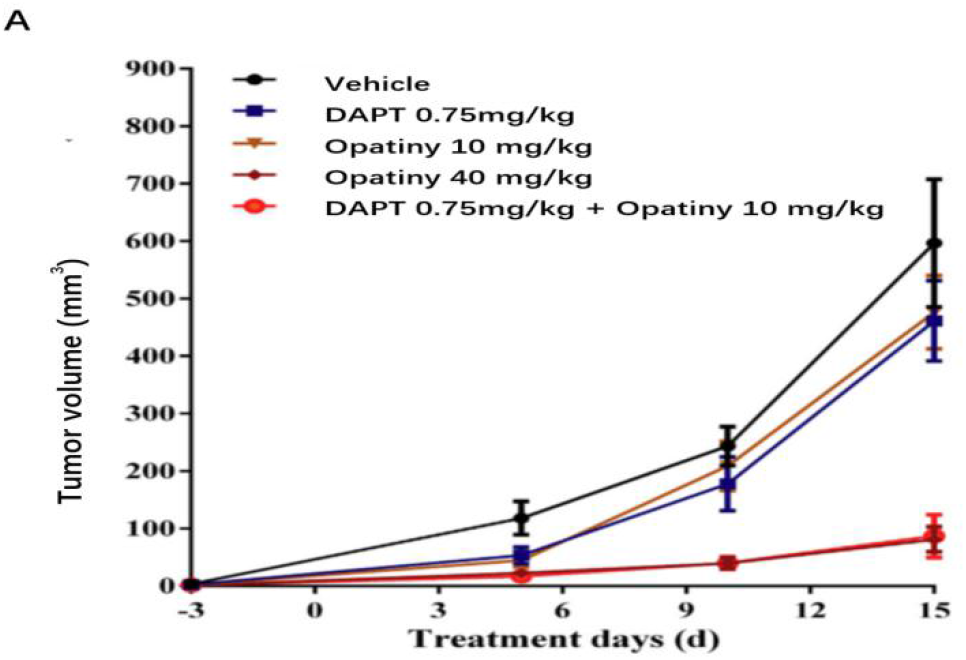
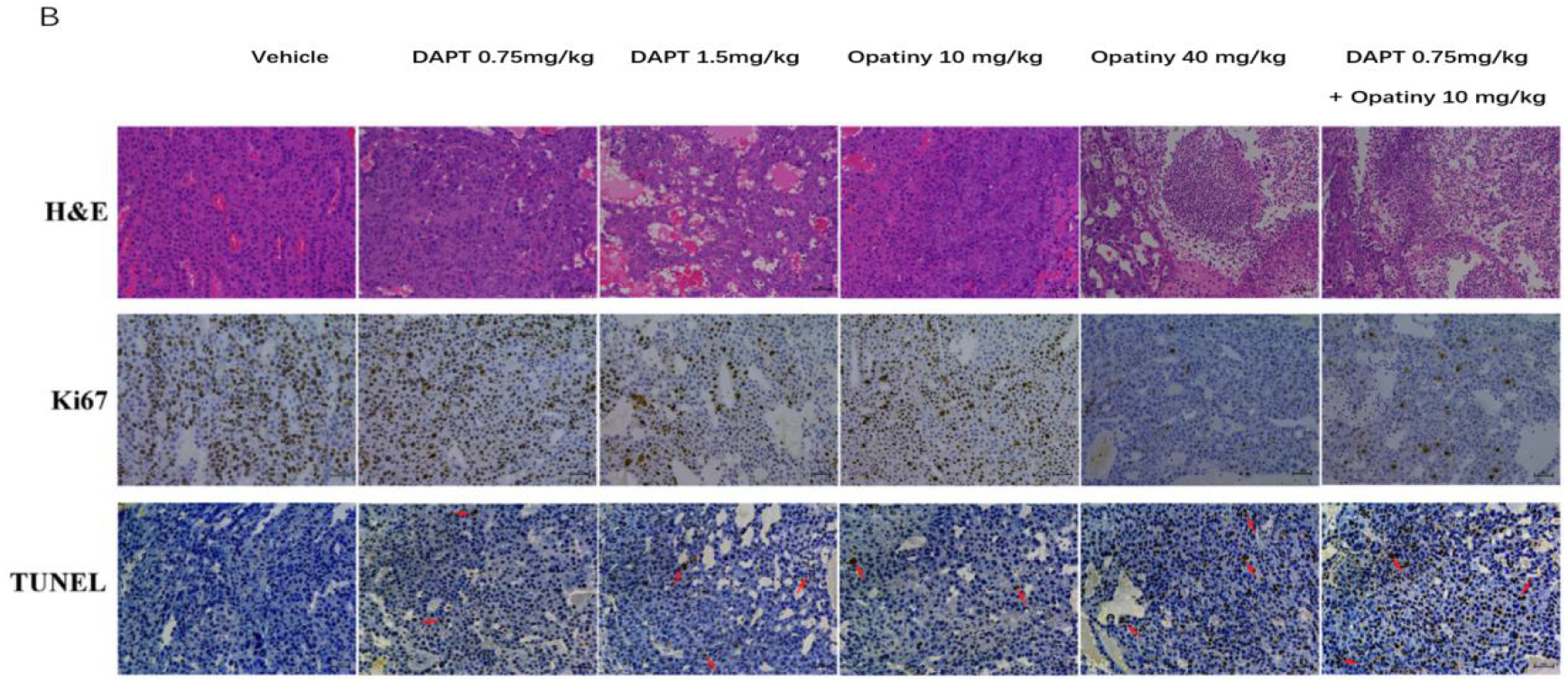
DAPT enhances the antitumor activity of olaparib in vivo (A) Growth curve of liver tumor in OVcar3 orthotopic transplanted mouse model. The data are expressed as mean ± SD. * p <0.05. (B) Representative hematoxylin and eosin (H & E) stained images of liver tumors in situ after DAPT and olaparib treatment. (C, D) Representative immunohistochemical staining of proliferation marker ki67 and apoptosis marker TUNEL (D) in tumors treated with DAPT and olaparib. Scale bar: 50 μm.

In order to examine the effect of DAPT on the antitumor activity of olaparib in vitro, we harvested tumor tissues after sacrificed mice to make paraffin sections, performed immunohistochemical staining, and analyzed apoptosis by TNUNNEL. The results showed that the combination of DAPT and olaparib can significantly induce tumor cell apoptosis (Figure 4B), reduce the number of ki67-positive cells (Figure 4C), and increase cancer tissue compared with low/high dose olaparib treatment. The number of TUNEL positive (Figure 4D), indicating that DAPT can significantly enhance the anti-ovarian cancer activity of olaparib in vivo.

### 3.5 DAPT inhibits the synthesis of Jagged1 in ovarian cancer cells

In order to detect the effect of DAPT on the expression of Jagged1 in ovarian cancer cells, we used 2 μM and 4 μM DAPT to treat ovarian cancer cells for 24 hours and then performed Western blot analysis and RT-PCR detection. The results showed that after DAPT treatment, Jagged1 protein expression levels and mRNA transcription levels were significantly reduced (Figure 5A, B, and C), and were time- and dose-dependent.

**Figure 5.**
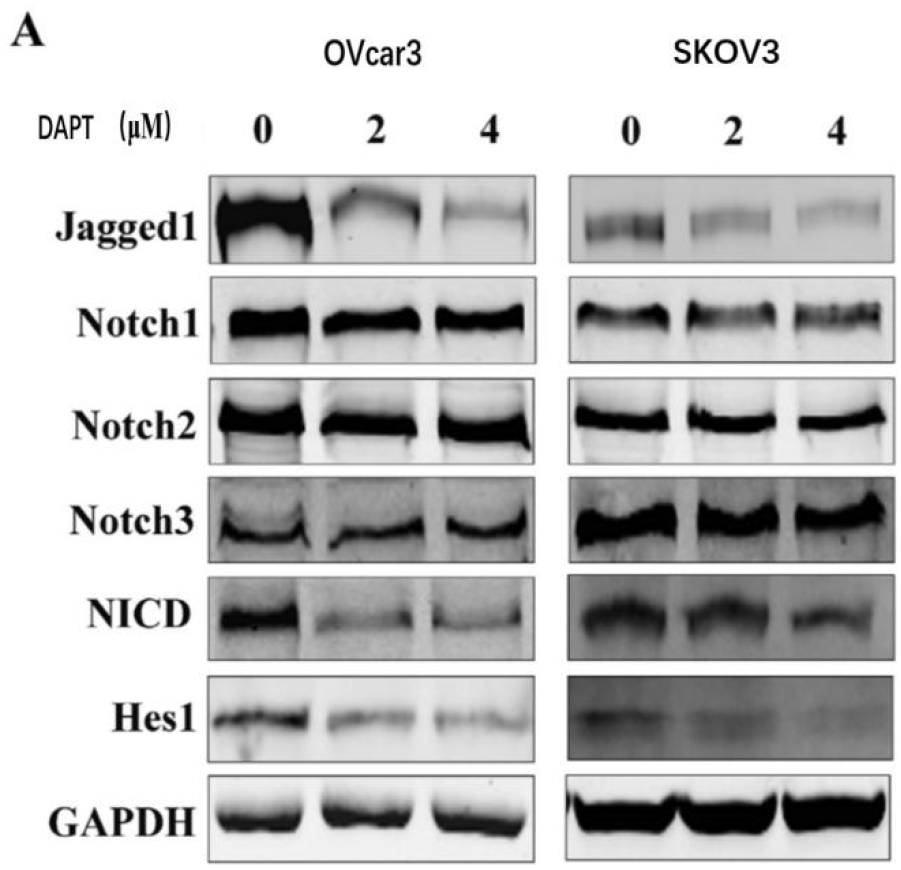
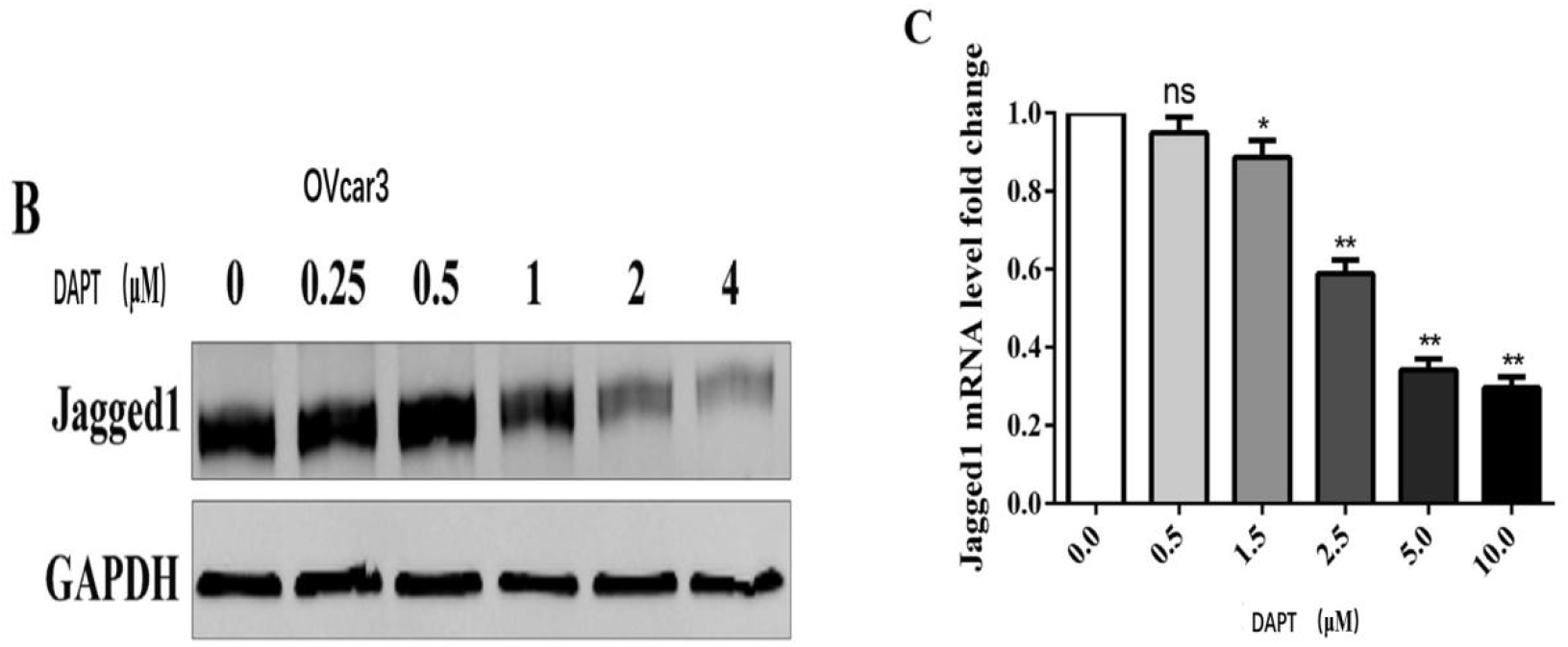
DAPT inhibits Jagged1 expression (A) The expression of related proteins in ovarian cancer cells after DAPT treatment, GAPDH was used as a control. (B) Western blot analysis of Notch ligand Jagged1 in OVcar3 cells after 24 hours of DAPT treatment (C) RT-qPCR analysis of Notch ligand Jagged1 in OVcar3 cells after 24 hours of DAPT treatment. Compared with the control group, * P <0.05, ** P <0.01.

### 3.6 DAPT inhibits β-catenin / Jagged1 signal transduction

Notch signal is the intersection of many important signaling pathways, and it is abnormally activated in many malignant tumors including liver cancer, gastric cancer, ovarian cancer, and colon cancer, and is closely related to the occurrence and development of tumors^[19–20]^. Its downstream signal Jagged1 is overexpression in 57% of human ovaries cancer, recent studies have shown that Jagged1 is a Wnt / TCF target gene, and targeted therapy against Jagged1 can prevent the development of tumors in a mouse model of primary liver cancer^[21–22]^.

In order to detect the antitumor target of DAPT and the correlation between DAPT antitumor activity and Jagged1, we used shJagged1 and negative control cDNA (siNC) to transfect ovarian cancer cells, and the cell viability was detected by CCK8 24 hours after transfection, and the ovarian cancer cells with Jagged1 knockdown were treated with/without 4 μM DAPT for 4 h and the cell viability was detected. The results showed that compared with the negative control group, the cell viability was significantly suppressed after Jagged1 knockdown, but there was no significant difference in cell viability after DAPT treatment, indicating that DAPT inhibits cell viability through Jagged1 and induces apoptosis.

In order to further detect the anti-tumor target of DAPT and the correlation of DAPT anti-tumor activity with β-catenin / Jagged1, we used shβ-catenin and negative control siRNA (siNC) to transfect ovarian cancer cells, treated with or without DAPT (4 μM) for 2 hours after transfection, and then detected the expression of related proteins by Western-blot.. The results showed that compared with the negative control group, after using shRNA (shβ-catenin) to knock down β-catenin expression, Jagged1 expression was significantly down-regulated, while promoting DAPT-mediated Jagged1 inhibition (Figure 6C), while in Jagged1 knockdown cells There is no change in β-catenin expression (Figure 6D), suggesting that β-catenin is the upstream signal of Jagged1, Jagged1 is the downstream target of β-catenin, and DAPT can down-regulate the expression of Jagged1 by inhibiting β-catenin.

**Figure 6.**
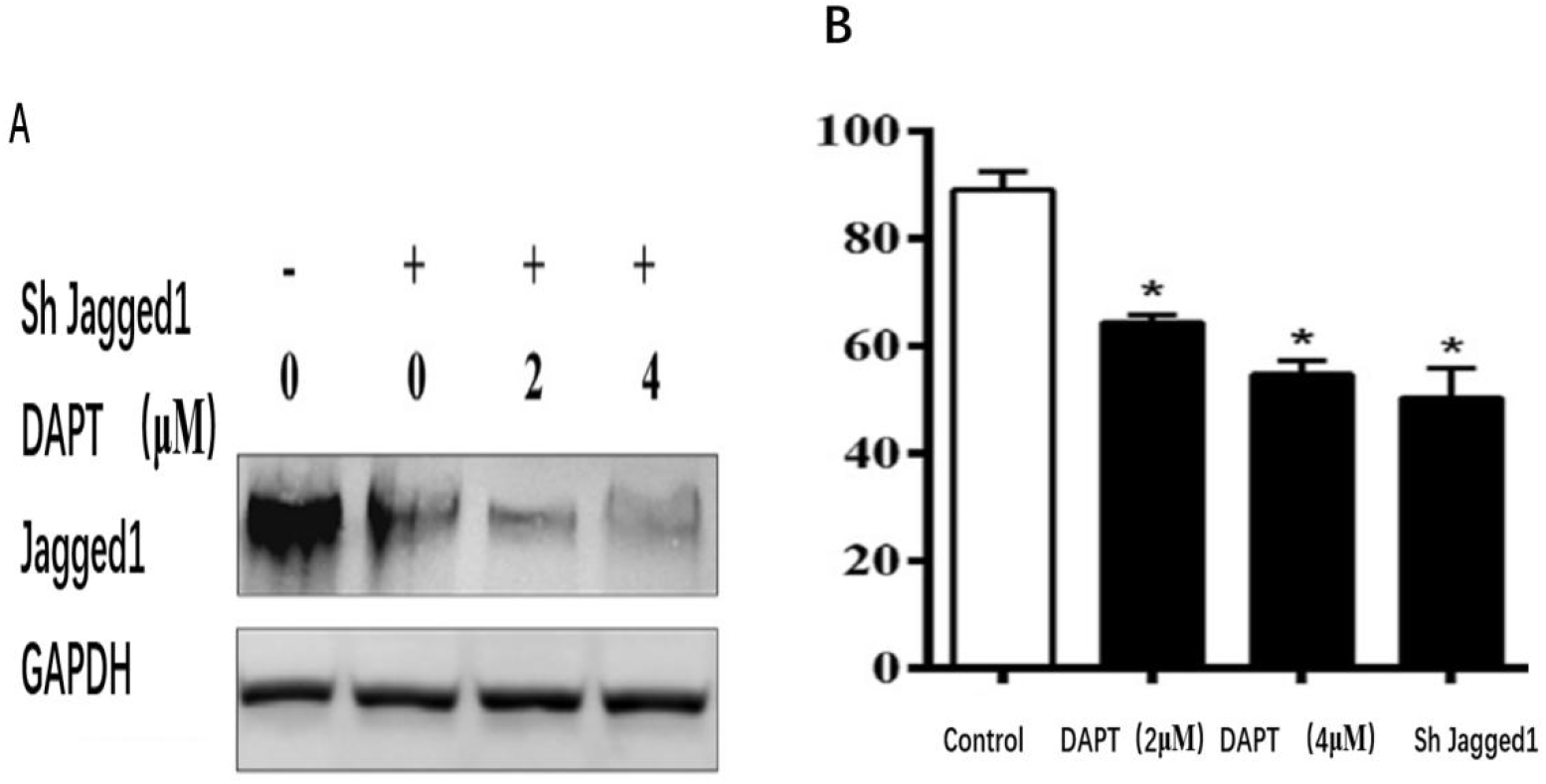
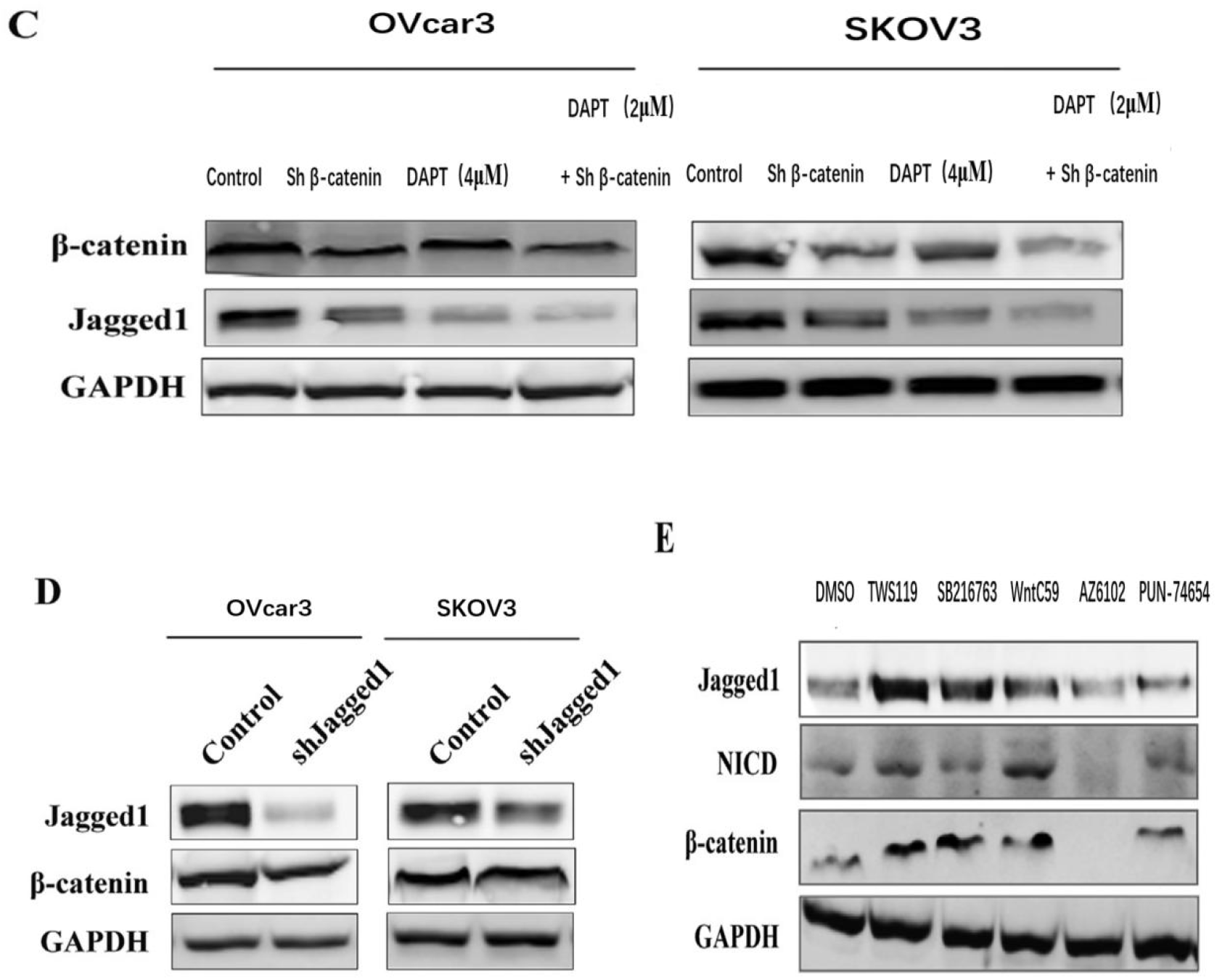
DAPT inhibits ovarian cancer cell proliferation through β-catenin / Jagged1 (A) OVcar3 cells over-expressed with Jagged1 or shRNA knockdown treated with DAPT were used to analyze the expression of related proteins by Western blotting. (B) Activity of OVcar3 cells after overexpression of DAPT or Jagged1 or shRNA knockdown treatment (C) Western blot images of Jagged1 in OVcar3 and SKOV3 cells after β-catenin knockdown. (D) Western blot analysis of β-catenin in OVcar3 and SKOV3 cells after Jagged1 knockdown. (E) Western blot analysis of OVcar3 cells after 24 hours of treatment with Wnt activator (25 ng · ml-1 SB216763), GSK3β inhibitor (2μM TWS119 and 3μM S8178) and Wnt inhibitor (10μM FH535 and 25μM ICG001).

To further confirm that Jagged1 is a downstream target of β-catenin, we used Wnt signaling inhibitors or Wnt activators to treat untransfected ovarian cancer cells, and then detected the expression of related proteins by Western-blot. The results showed that after using Wnt signaling inhibitors FH535 and ICG-001 to inhibit β-catenin expression, Jagged1 expression was significantly down-regulated, while Wnt activators SB216763, TWS119 and S8178 could significantly upregulate Jagged1 expression (Figure 6E), indicating that ovarian cancer Jagged1 is the downstream target of β-catenin, and Jagged1 may be the intermediate linking molecule between Notch and Wnt / β-catenin.

### 3.7 DAPT induces β-catenin degradation in a proteasome-dependent manner

β -catenin is the main downstream effector of the classic Wnt signaling pathway, mainly present in the cytoplasm. The cytoplasmic level is strictly controlled by the complex composed by adenomatous polyposis (APC), AXIN and glycogen synthase kinase 3β (GSK3β), and the complex induces its ubiquitination and proteasome-mediated degradation by phosphorylating serine on amino acid residue 37 (Ser37) or threonine on amino acid residue 41 (Thr41) of β-catenin^[23–24]^. Wnt signal transduction can induce β-catenin to enter the nucleus, regulate specific oncogenes, including c-MYC, cyclin D1 and survivin, and activate transcription factors together with the transcription enhancement factor / T cytokine (LEF / TCF) family to participate the occurrence and development of tumors^[25–26]^.

In order to detect the effect of DAPT on Wnt / β-catenin signal transduction and its downstream targets, we used DAPT to treat ovarian cancer for 4h and then detected the transcription of related genes and the expression of related proteins by Western blot and PCR. The results show that DAPT can significantly reduce the expression levels of Wnt / β-catenin target genes LEF1, survivin, AXIN-2 and c-Myc in OVcar3 cells in a dose-dependent manner (Figure 7A), and down-regulate the protein level of Wnt / β-catenin target genes, and showed a dose and time dependence (Figure 7B), indicating that DAPT can inhibit Wnt / β-catenin signaling by targeting β-catenin.

**Figure 7.**
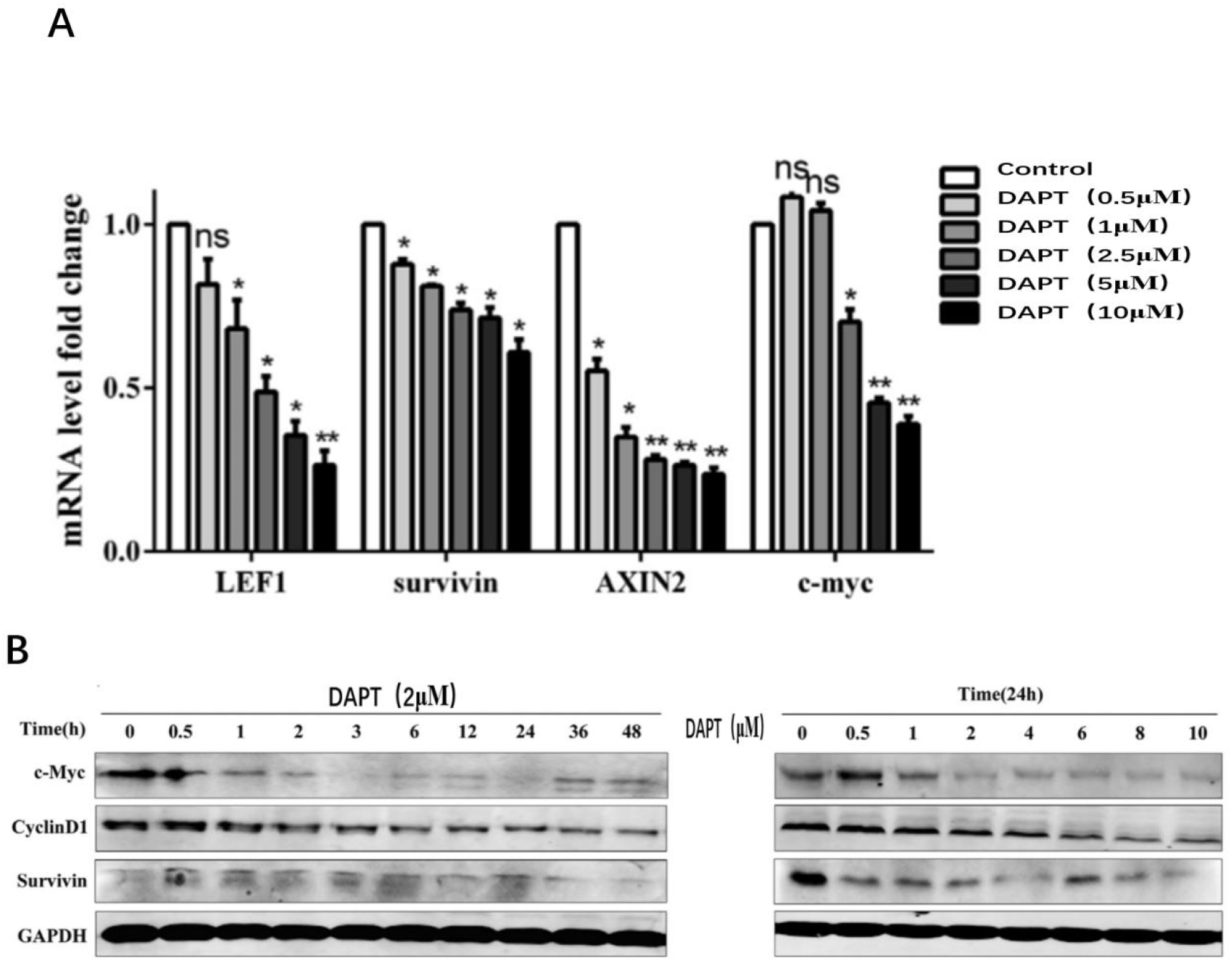
DAPT inhibits Wnt transcription activity and reduces Wnt target gene expression. (A) Levels of Wnt / β-catenin target genes Lef1, survivin, Axin2 and c-myc in OVcar3 cells after DAPT treatment for 24 hours. (B) Western blot analysis of DAPT-treated OVcar3 cells at different time points.

In order to further detect the potential mechanism of DAPT inhibition of β-catenin, we used Western blot and immunofluorescence to detect the nuclear expression level and cytoplasmic level of β-catenin after treatment with DAPT (4 μM), while Wnt / β-catenin agonist or proteasome inhibitors were used to treat with ovarian cancer cells, and then Western blot was used to detect the expression level of β-catenin. The results showed that nuclear β-catenin levels were significantly reduced after DAPT treatment, while cytoplasmic levels were not affected (Figure 8A-B). Total β-catenin and active β-catenin (ABC) in cell lysates were significantly reduced, indicating that DAPT down-regulated the overall level of β-catenin and inhibited nuclear β-catenin and ABC levels (Figure 8C).

**Figure 8.**
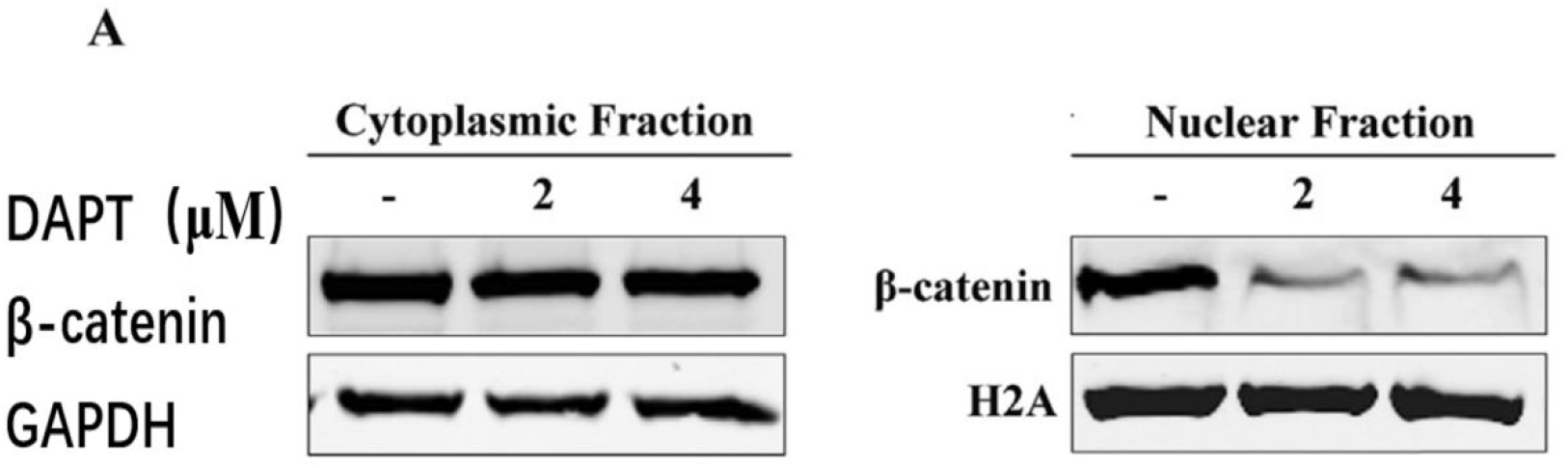
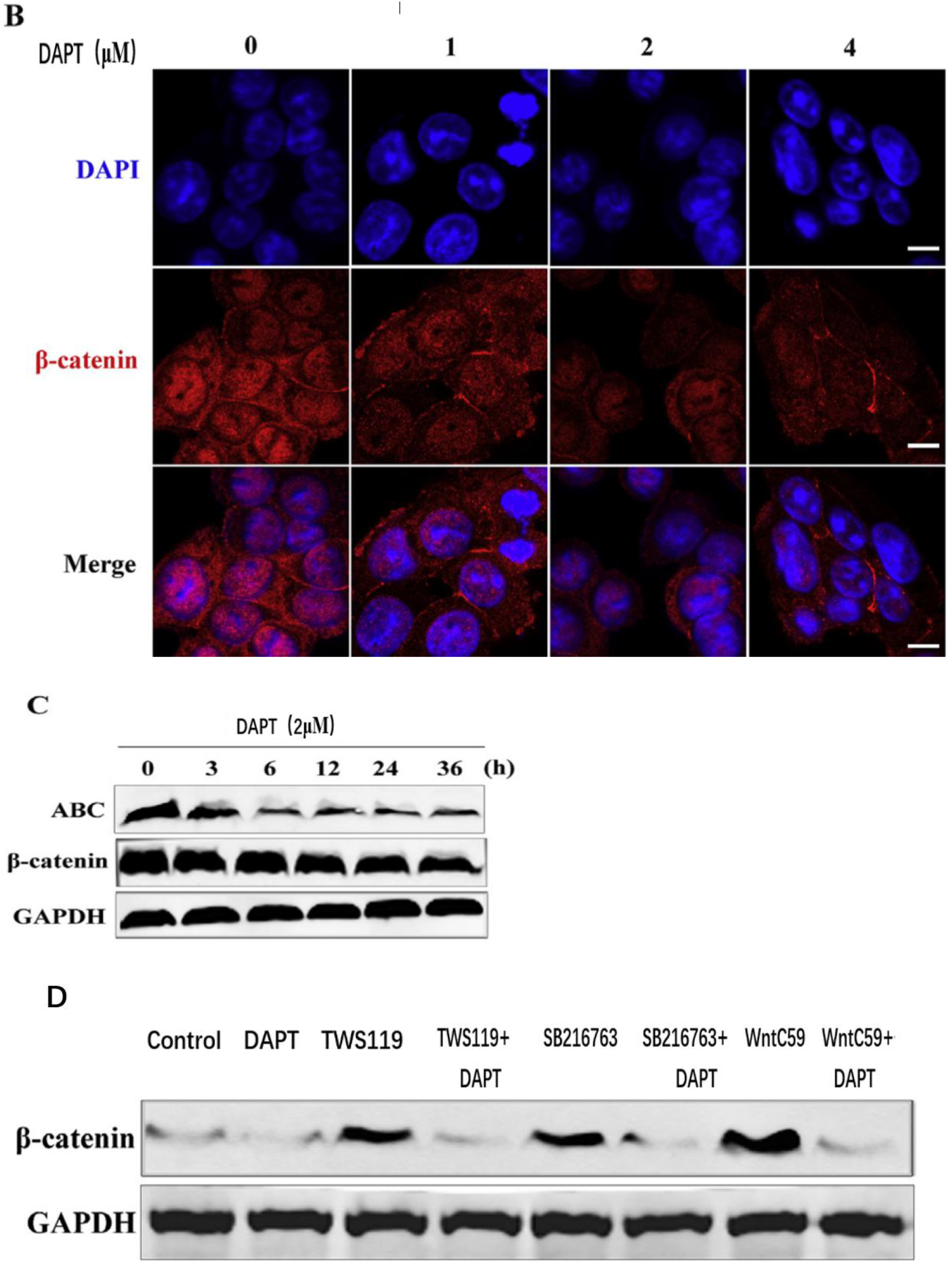
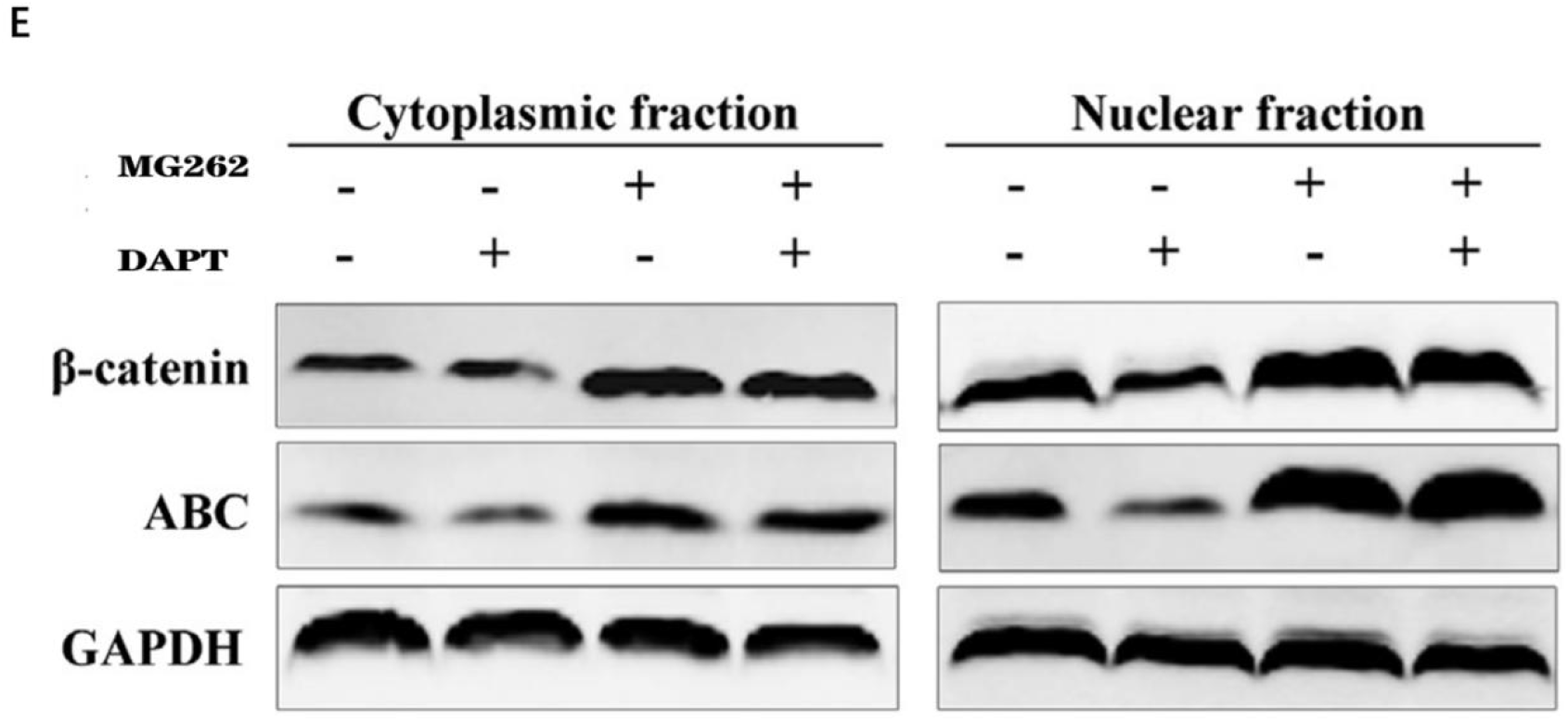
DAPT reduces Wnt-mediated β-catenin nuclear accumulation in OVcar3 cells (A) Western blot analysis of cell lysates after treating OVcar3 cells with different concentrations of DAPT for 24 hours (B) Representative immunofluorescence images of OVcar3 cells after 24 hours of treatment with DAPT. Scale bar = 20 mm. (C) Western blot analysis of OVcar3 cells at different time points after treatment with DAPT (2 μM). (D) RT-qPCR analysis of OVcar3 cells after treatment with DAPT (2μM) for a specified period of time. (E) Western blot analysis of OVcar3 cells treated with DAPT alone or in combination with 25 ng·ml-1 SB216763, 2 μM TWS119 or 3 mM S8178 after treatment for 24 hours (F) OVcar3 cells were treated with 2 μM DAPT for 16 hours, and then cultured with or without the 10 μM proteasome inhibitor MG132 for 12 h before Western blot analysis.

In addition, DAPT can significantly reduce the total β-catenin level in OVcar3 cells after SB216763-, TWS119 and S8178 treatment (Figure 8E), while in the presence of proteasome inhibitor MG132, DAPT does not change the β-catenin level (Figure 8F), indicating, DAPT induces β-catenin degradation in a proteasome-dependent manner and thus inhibits Wnt / β-catenin-mediated translation activity.

In addition, the immunohistochemical staining in this study further confirmed the reduction of β-catenin, ABC and Jagged1 protein expression after DAPT treatment (Figure 9A-B). The results of immunohistochemistry and Western blot analysis indicate that DAPT can inhibit the progression of ovarian cancer in vivo and in vivo by targeting the β-catenin / Jagged1 cascade.

**Figure 9.**
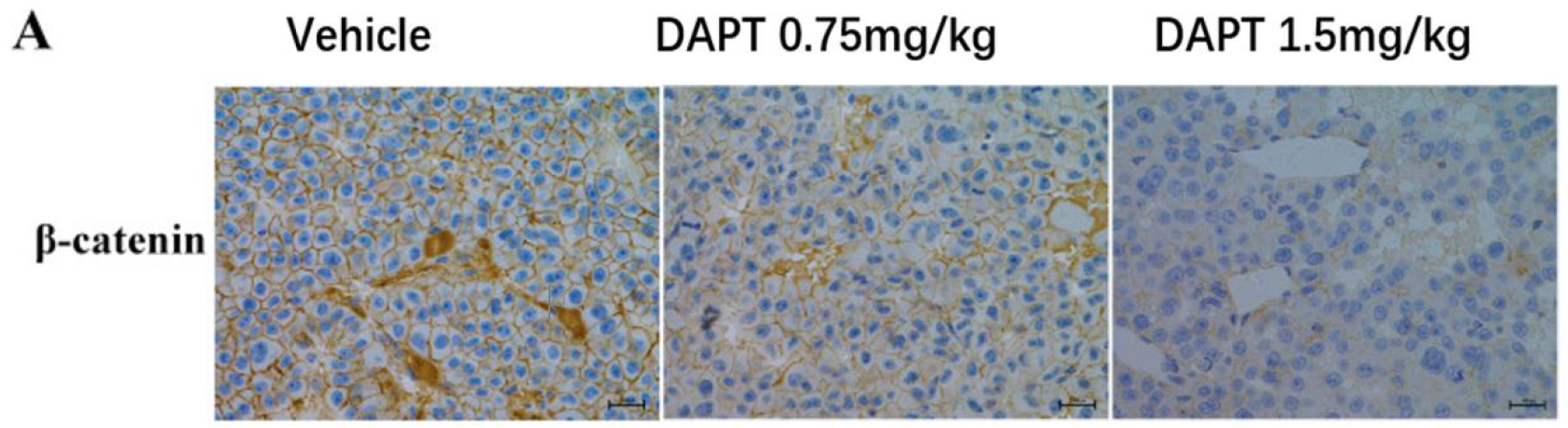
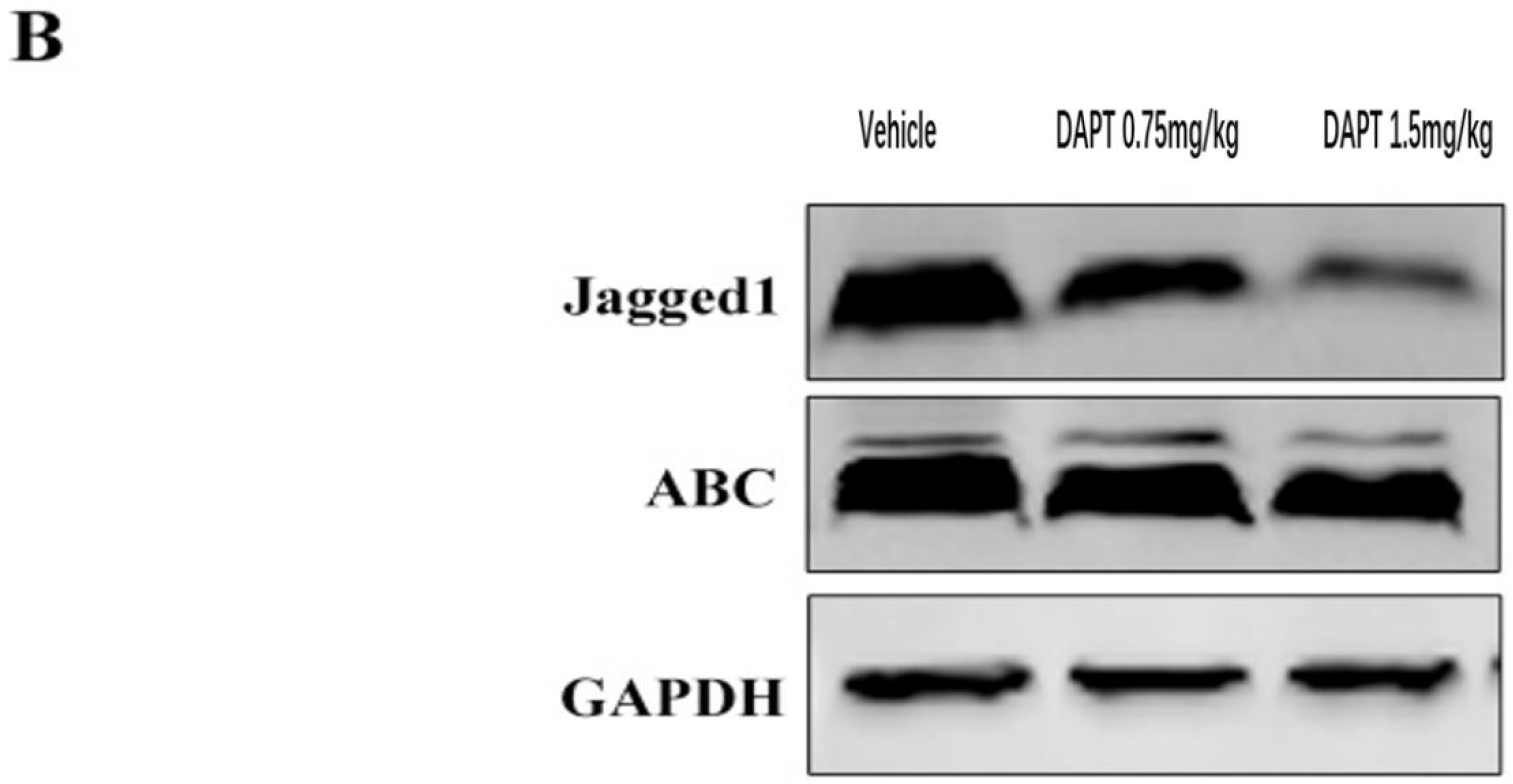
DAPT inhibits the expression of β-catenin and Jagged1 in an orthotopic mouse tumor model. (A) Representative immunohistochemical staining of β-catenin in tumor tissue after DAPT treatment. Scale bar: 50 μm. (B) Western blot analysis of Jagged1 and active β-catenin (ABC) in tumor samples treated with DAPT at different concentrations. Scale bar: 50 μm.

## 4 Discussion

Ovarian cancer is one of the common malignant tumors in gynecology, endangering the physical and mental health of female patients^[27]^. Due to its concealed characteristics, it is often found in the middle and late stages, and the first-line chemotherapy drugs for advanced ovarian cancer including apatinib have limited clinical benefits due to the acquisition of chemotherapy resistance^[28]^. In this study, DAPT was used to treat ovarian cancer cells and xenograft mouse models. The results showed that DAPT treatment can significantly inhibit the progress of ovarian cancer in vivo and in vitro. In addition; compared with low-dose apatinib treatment, the combination of DAPT and apatinib can significantly inhibit the progression of ovarian cancer in vivo and in vivo, and shows similar effects as high-dose apatinib treatment, indicating that DAPT can enhance apatinib The anti-tumor activity of nipa is better than that of apatinib alone.

Notch1 / Jagged1 and Wnt / β-catenin signaling play an important role in the development of ovarian cancer. The Notch gene was discovered in Drosophila melanogaster in 1919^[29]^. It was originally discovered that the deletion of this gene can lead to notches in the wings of Drosophila, and hence its name, participating in the maintenance of cell homeostasis is a relatively conservative signaling pathway. In mammals, the Notch signaling pathway includes four receptors and five ligands, of which Jagged1 is an important downstream molecule of the signaling pathway, and its expression is increased in many solid tumors such as lung cancer, cervical cancer, colon cancer and kidney cancer,and through the excessive activation of Notch1 / Jagged1 in malignant tumor tissue to promote cell proliferation and invasion^[30–32]^. The results of this study show that DAPT treatment can significantly inhibit the expression and transcription of Jagged1, and gene knockout experiments further indicate that Jagged1 may be a potential downstream target of DAPT tumor suppressor activity.

Recent studies have shown that Jagged1 is a potential downstream target gene of Wnt / TCF^[33]^. Targeted therapy for Jagged1 can prevent the development of tumors in a mouse model of primary ovarian cancer, while studies in mammalian cells have shown that Notch1 can bind and Antagonizes β-catenin, and β-catenin can activate Jagged1 transcription, leading to excessive activation of Notch in colorectal cancer and breast cancer, indicating that there is an interaction between Wnt and Notch signaling, and β-catenin plays a transit role in it^[34–35]^. Wnt / β-catenin signaling is an evolutionarily conserved pathway that plays an important role in protecting ovarian health and promoting homeostasis in the ovary. Recent studies have shown that abnormal activation of the Wnt / β-catenin pathway is closely related to the development of various malignant tumors included in ovarian cancer ^[36]^.In this study, gene knockout was used to detect the interaction between Wnt / β-catenin and Jagged1. The results showed that there was no significant change in β-catenin expression when Jagged1 was knocked down, and β-catenin knockdown could significantly reduce Jagged1 expression and transcription. In addition, DAPT can significantly promote β-catenin knockdown-induced Jagged1 inhibition, indicating that Jagged1 is a downstream target of β-catenin, and DAPT exerts antitumor activity through β-catenin / Jagged1. In order to detect the inhibitory effect of DAPT on β-catenin and its potential mechanism, this study further used Wnt / β-catenin agonists or proteasome inhibitors in combination with DAPT to treat ovarian cancer cells. The results show that DAPT can lead to a significant reduction in the nuclear accumulation of β-catenin in ovarian cancer cells and a significant reduction in ABC, thereby inhibiting the expression of Wnt target genes Lef1, Survivin, Axin2 and c-Myc and related protein levels, while in the presence of the proteasome inhibitor, DAPT does not change the level of β-catenin, indicating that DAPT induces β-catenin degradation in a proteasome-dependent manner and thus inhibits Wnt/β-catenin-mediated translation activity.

In conclusion, our research results show that DAPT can inhibit cell proliferation, induce apoptosis, inhibit tumor development in vivo through β-catenin / Jagged1, and play the sensitizing effect of olaparib on ovarian cancer activity. Our research will provide a theoretical basis for DAPT as a potential drug for ovarian cancer treatment or olaparib sensitizer for clinical adjuvant treatment of ovarian cancer patients.

